# H5N1 influenza binding and cell entry via human class II MHC, and blocking by cross-reactive antibodies

**DOI:** 10.64898/2026.07.22.739677

**Authors:** Taylor Pursell, Artem Mikelov, Oliver F. Wirz, Jordan T. Ort, Shuk Hang Li, Reilly K. Atkinson, Jefferson J. S. Santos, Jiabao Zhong, Shilpa A. Joshi, Jumana Afaghani, Xiaorui Han, Emily Haraguchi, Ramona A. Hoh, Ji-Yeun Lee, Brandon Lam, Alexander Stanford, Andrew T. DeLaitsch, Jackson Schuetz, Katharina Röltgen, Donna Smith, Brian Ha, Sean Van Slyck, Claus U. Niemann, Scott E. Hensley, Scott D. Boyd

## Abstract

Highly pathogenic avian influenza (HPAI) H5N1 clade 2.3.4.4b viruses are currently responsible for a multi-species outbreak affecting wild birds, poultry, numerous mammalian species, and humans. Influenza A viruses typically initiate infection through binding to sialic acid, although select bat and human influenza viruses can also exploit class II major histocompatibility complex (MHC-II) molecules for cell entry. Here we show that emerging H5N1 clade 2.3.4.4b viruses, but not historical H5 lineages, bind human MHC-II HLA-DR and mediate sialic acid-independent cell entry. Hemagglutinin binding to primary human immune cells varies with MHC-II expression and is further shaped by HLA-DR allelic variation, identifying host genetic determinants that may influence susceptibility to infection. Mammalian-adaptive substitutions within the hemagglutinin sialic acid receptor-binding domain reduce MHC-II binding, suggesting this interaction is remodeled during clade 2.3.4.4b H5 adaptation to a human host. Lastly, cross-reactive monoclonal antibodies isolated from clade 2.3.4.4b H5-naive humans can block the hemagglutinin–MHC-II interaction. These findings identify a previously unrecognized receptor pathway in contemporary H5N1 viruses and reveal that both human genetic variation and pre-existing humoral immunity can modulate this interaction, with implications for host range, cellular tropism, spillover risk, and therapeutic intervention.

---

Influenza A viruses (IAV) have long been understood to utilize cell-surface sialic acid moieties as the primary determinant for host-cell recognition and entry^1^. The viral hemagglutinin (HA) protein mediates initial binding to sialic acid receptors on respiratory epithelial cells, initiating a cascade of conformational changes that enables fusion with endosomal membranes^2^. This receptor-binding paradigm has shaped our understanding of IAV species and tissue tropism, host adaptation, and pandemic potential for over a century. However, recent evidence has challenged the sufficiency of the sialic acid-only model. Bat-derived IAVs utilize MHC-II molecules exclusively as functional receptors^3^, while selected historical human, avian, and swine IAV strains, including human H2 influenza from the 1957 to1968 pandemic possess MHC-II binding capacity, typically for HLA-DR homologs^4–6^. These discoveries suggest that MHC-II engagement is an ancient and broadly distributed influenza viral entry mechanism, potentially relevant to understanding contemporary strains.

Since its emergence in 2013, HPAI H5N1 clade 2.3.4.4b viruses have demonstrated unprecedented geographic spread and cross-species transmission capacity, affecting >100 countries and infecting diverse mammalian species including seals, mink, and cattle^7^. Notably, adaptation to mammalian hosts appears to involve specific amino acid changes within the HA1 receptor-binding domain^8^. Whether these adaptations include MHC-II engagement remains unknown. Here, we demonstrate clade 2.3.4.4b H5 strains exhibit dual receptor specificity, utilizing MHC-II HLA-DR as well as sialic acid entry receptors. This capacity is conserved across recent circulating viral isolates, is functionally relevant for cell entry in primary airway epithelial cells and macrophages, and is affected by HLA-DR genotypes and pre-existing influenza-specific antibodies. Our findings broaden the understanding of IAV receptor utilization and suggest that H5N1 strains and other influenza viruses capable of MHCII-dependent entry may pose enhanced zoonotic risk, warranting inclusion in pandemic preparedness surveillance and vaccine development strategies.

## Human MHC-II binding is a feature of highly pathogenic avian influenza H5N1 clade 2.3.4.4b strains

Our investigations began with a serendipitous observation while attempting to use the hemagglutinin of clade 2.3.4.4b H5 influenza for flow cytometric sorting of H5-specific B cells from primary human blood or spleen samples. We labeled cells with a Y98F mutant (H3 numbering system) prototypic clade 2.3.4.4b H5 HA (A/American Wigeon/South Carolina/22-000345-001/2021) that should lack sialic acid binding, but that nonetheless bound to a large fraction of total human B cells tested (up to 22%, Fig. 1a upper left panel). These binding frequencies are inconsistent with true BCR-mediated binding by antigen-specific memory B cells, which are typically found at much lower frequencies for most viral antigens in human blood or spleen (Fig. 1a). Given the broad host range of the emergent H5N1 clade 2.3.4.4b viruses, and recent reports of dual sialic acid and MHC-II receptor binding by H2 and H3 human influenza isolates^4,5^, we hypothesized that the emergent H5N1 clade 2.3.4.4b HA might also exhibit MHC-II binding.

**Figure 1.**
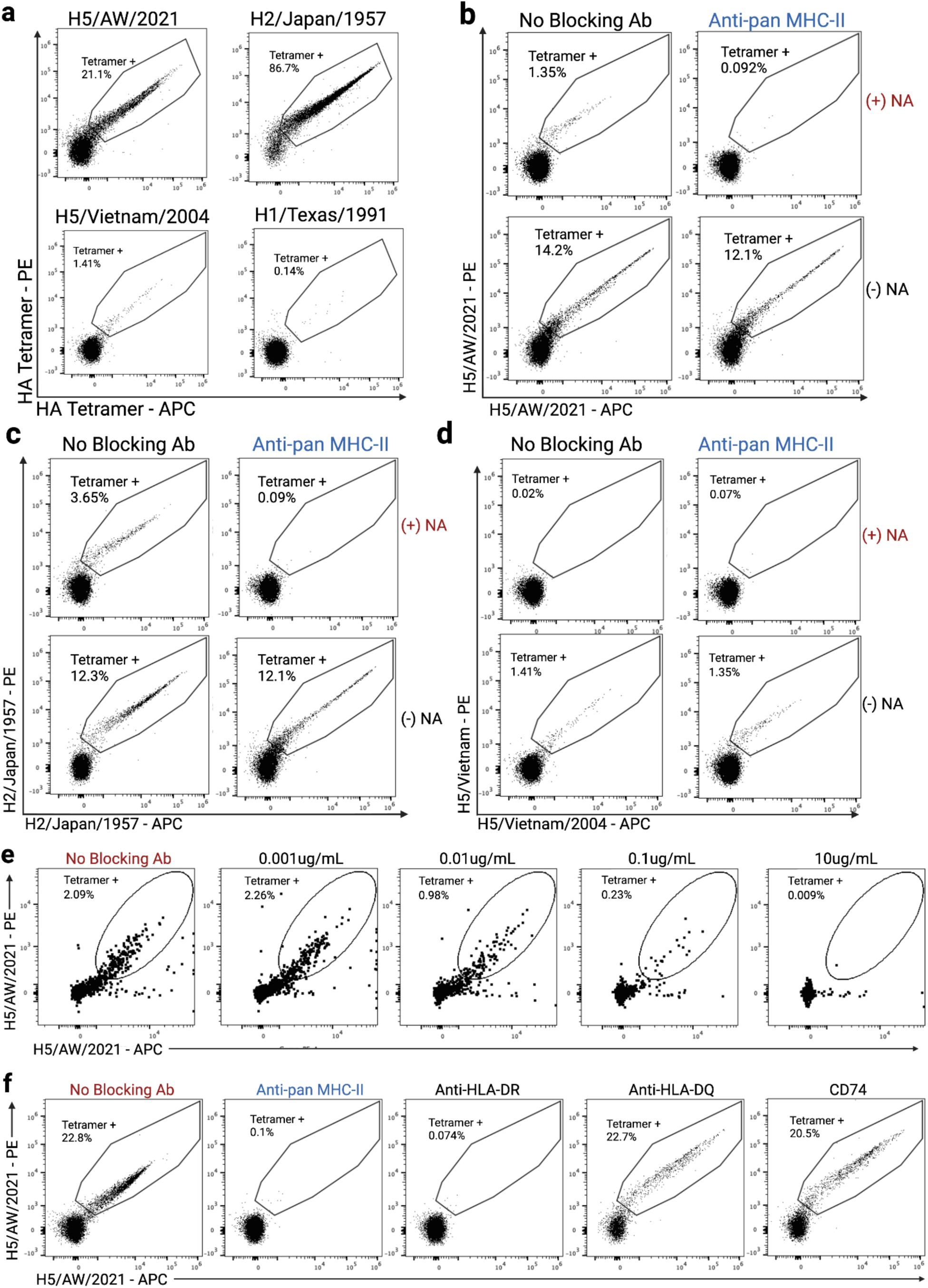
Clade 2.3.4.4b HA binds human MHC-II. **a,** Representative flow cytometry plots of pooled human B cells stained with HA tetramers from clade 2.3.4.4b H5 (A/American Wigeon/South Carolina/22-000345-001/2020-Y98F), H2 (A/Japan/305/1957-Y98F), clade 1 H5 (A/Vietnam/1203/2004-Y98F), and H1 (A/Texas/36/1991-Y98F). **b-d,** Representative flow cytometry plots of pooled human B cells stained with **b,** clade 2.3.4.4b H5/AW/2021-Y98F tetramer **c,** H2/japan/1957-Y98F and **d,** clade 1 H5/Vietnam/2004-Y98F tetramer following no treatment, anti-MHC-II blocking (blue), neuraminidase treatment (red, NA), or neuraminidase treatment combined with anti-MHC-II blocking. **e,** Titration of anti-MHC-II blocking antibody on neuraminidase-treated pooled human B cells stained with the clade 2.3.4.4b H5/AW/2021-Y98F tetramer. **f**, Binding of the clade 2.3.4.4b H5/AW/2021-Y98F tetramer to neuraminidase-treated pooled human B cells following blockade with pan-MHC-II, HLA-DR, HLA-DQ, or CD74 antibodies.

To test this hypothesis, we used human peripheral blood mononuclear cells (PBMCs), which are a mixture of monocytes, B cells, T cells, NK cells, and small fractions of other cell types; of these, most display abundant sialic acid on their surface, while monocytes and subsets of B cells are the most prominent populations with elevated MHC-II expression. We used fluorescently tagged streptavidin HA tetramers to screen for binding of individual HAs to B cells from pooled human PBMCs. We either treated cells with neuraminidase to remove sialic acid or left cells untreated before staining with HA tetramers of select influenza strains containing the Y98F mutation: A/Vietnam/1203/2004-Y98F (clade 1, H5/Vietnam/2004), A/American Wigeon/South Carolina/22-000345-001/2021-Y98F (clade 2.3.4.4b; H5/AW/2021-Y98F), A/Japan/305/1957 (H2/Japan/1957-Y98F, which binds both sialic acid and HLA-DR^5^), and A/Texas/1991-Y98F (H1). Untreated cells showed low binding of H1-Y98F tetramers, consistent with binding to antigen specific B-cells, but H5/AW/2021-Y98F, H2/Japan/1957-Y98F and, to a lesser extent, H5/Vietnam/2004-Y98F HAs demonstrated significantly more binding to PBMCs (Fig. 1a). Clade 2.3.4.4b H5/AW/2021-Y98F (Fig. 1b) and H2/Japan/1957-Y98F (Fig. 1c) tetramers showed residual elevated binding to the B cells after neuraminidase treatment, but clade 1 H5/Vietnam/2004-Y98F (Fig. 1d) tetramers binding decreased to below H1-Y98F levels. Only in the presence of both neuraminidase treatment and preblocking of cells with an anti-MHC-II antibody did we see binding similar to, or less than, H1-Y98F frequencies (Fig. 1b & 1c, upper right panel). To validate specificity, we titrated the blocking anti-MHC-II antibody and saw a negative correlation between concentration of blocking antibody and the number of tetramer positive cells detected (Fig. 1e). We then tested blocking antibodies specific for the different MHC-II subtypes (HLA-DR, HLA-DQ, and CD74), and found that only HLA-DR blockade with an antibody shown to block HA-MHC-II binding in other influenza strains^3,5,6^ prevented H5/AW/2021-Y98F HA binding. The extent of blocking achieved with the pan-MHC-II antibody was similar to that of the anti-HLA-DR-specific monoclonal antibody, suggesting that HLA-DR binding accounts for most of the H5/AW/2021 HA binding (Fig. 1f).

To further evaluate whether HLA-DR expression was sufficient for H5/AW/2021 HA binding to cells, we transiently transfected HEK-293T cells, which lack HLA-DR expression, with plasmids encoding alpha and beta chains of human HLA-DR or a control HLA-DM receptor with C-terminal epitope tags. After 24 hours, cells were either left untreated or neuraminidase treated before being stained with fluorophore-conjugated antibodies specific for epitope tags on the HLA proteins and fluorophore-conjugated HA tetramers. Depletion of ɑ-2,3 sialic acid (SA) (Fig. 2a) and surface expression of constructs (Fig. 2b & 2c) were validated via flow cytometry. Neuraminidase treatment significantly decreased the percentage of clade 2.3.4.4b H5/AW/2021-Y98F tetramer positive cells (Supplemental Fig. 1). Transfection with HLA-DR increased binding by clade 2.3.4.4b H5/AW/2021-Y98F tetramers and H2/Japan/1958-Y98F but not by clade 1 H5/Vietnam/2004-Y98F or H1-Y98F controls (data not shown). The clade 2.3.4.4b H5/AW/2021-Y98F HA binding was inhibited by addition of an anti-HLA-DR monoclonal, similar to H2/Japan/1957-Y98F control (Fig. 2d & 2e).

**Figure 2.**
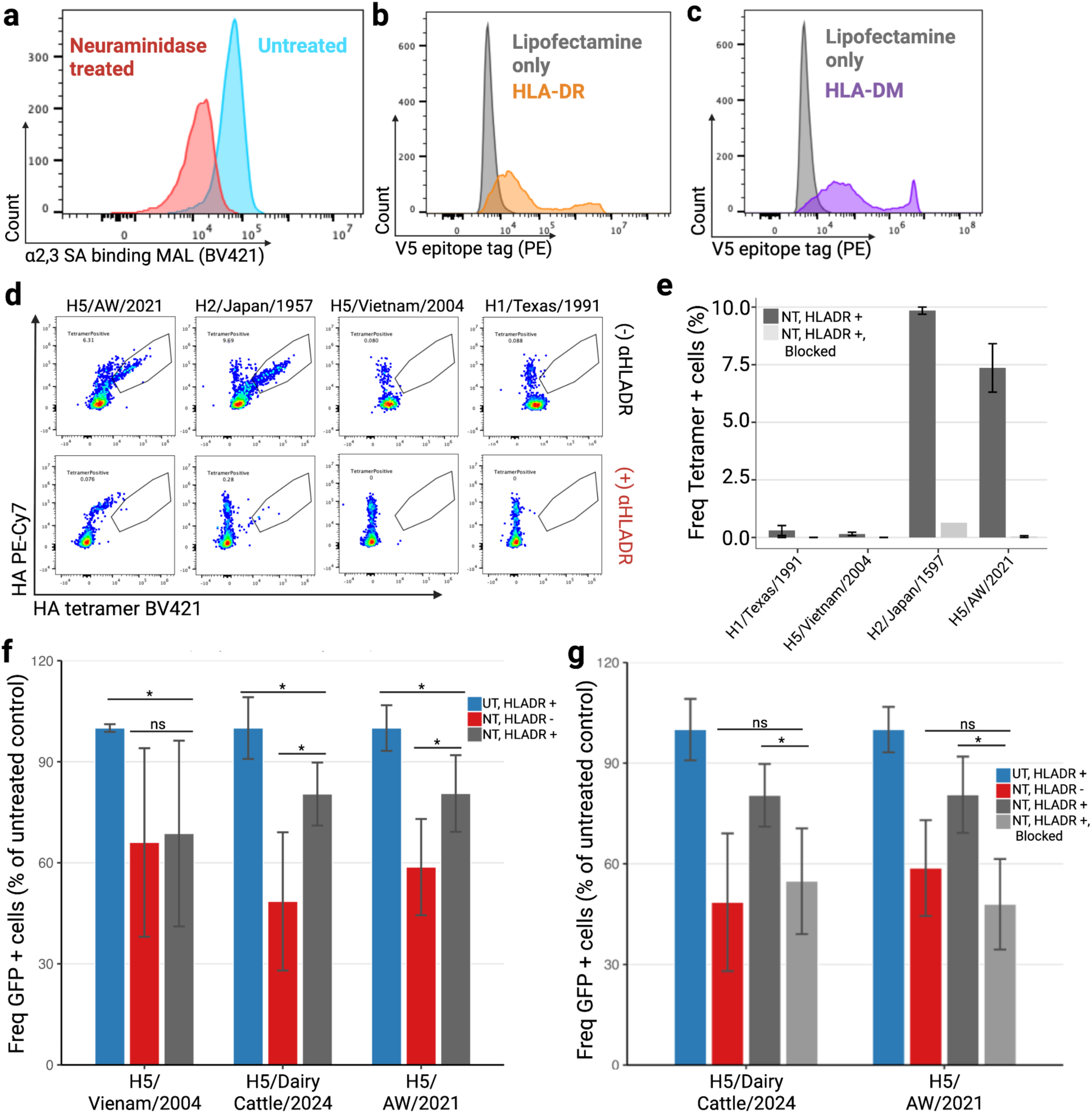
HPAI clade 2.3.4.4b H5N1 influenza strains can utilize HLA-DR to mediate entry. **a,** Histogram of ɑ-2,3 sialic acid binding lectin MAL II BV421 staining of untreated cells in blue and neuraminidase treated HEK293T cells in red. **b,** Histogram of HLADRB-V5tag PE staining of HLADR transfected cells (orange) and lipofectamine only control (grey). **c,** Histogram of HLADM-V5tag PE staining of HLADM transfected cells (purple) and lipofectamine only control (grey). **d,** Representative FC plots of live, HLADRA positive HEK293T cells either without (top row) or with anti-HLADR blocking (bottom row, red) gated on tetramer positive cells for cells for a panel of tetramers (from left to right H5/AW/2021-Y98F, H2/Japan/1957-Y98F, H5/Vietnam/2004-Y98F, or H1/Texas/1991-Y98F). **e,** Bar plot of frequency of tetramer positive, neuraminidase treated HEK293T cells either HLA-DR positive (dark grey) or HLA-DR positive with anti-HLAD-R blocking (light grey). Error bars represent standard deviation of two technical replicates (n=2). **f,** Bar plot of frequency of GFP positive HEK293T cells untreated and HLA-DR positive (blue), neuraminidase treated (red), or neuraminidase treated and HLA-DR positive (dark grey). Error bars represent standard deviation of three experimental replicates each with two technical replicates. * p< 0.05, n.s. indicates not significant. **g,** Bar plot of frequency of tetramer positive, neuraminidase treated HEK293T cells either lipofectamine alone (red), HLA-DR positive (dark grey) or HLA-DR positive with anti-HLADR blocking (light grey). Error bar represents the standard deviation of two experimental replicates (n=2).

## Human HLA-DR mediates sialic acid-independent viral cellular entry

To test whether human HLA-DR can contribute to cellular entry by HPAI clade 2.3.4.4b H5 influenza HA, we incubated with GFP reporter virus particles (RVPs) with HEK-293T cells either mock transfected or transiently expressing HLA-DR with or without SA depletion with neuraminidase treatment. RVPs for the following H5N1 strains were compared: clade 2.3.4.4b strain, A/American Wigeon/South Carolina/22-000345-001/2021 (H5/AW/2021), clade 2.3.4.4b A/dairy cattle/Texas/24-008749-003-original/2024 (H5/DairyCattle/2024), or control clade 1 A/Vietnam/1203/2004 (H5/Vietnam/2004). Under SA-depleted conditions without HLA-DR transfection, all strains demonstrated a significant reduction of the viral entry GFP reporter (48-60% reduction; adjusted p < 0.05, Fig. 2f). Both H5N1 clade 2.3.4.4b strains showed robust rescue of viral entry into cells that were transfected with HLA-DR (for H5/DairyCattle/2024, 40.7% increase, p = 0.008, 68% efficiency; for H5/AW/2021, 25.4% increase, p = 0.005, 50% efficiency), while control clade 1 H5/Vietnam/2004 strain showed no significant rescue (p = 0.558). To evaluate the requirement for HLA-DR-specific binding for this effect, HLA-DR-transfected cells were incubated with an anti-HLA-DR blocking antibody prior to inoculation with representative H5N1 clade 2.3.4.4b strains, H5/DairyCattle/2024 and H5/AW/2021. The addition of HLA-DR blocking monoclonal resulted in a significant decrease in entry positive cells to levels not significantly different from neuraminidase alone without HLA-DR transfection (Fig. 2g). Together these data support a role for human HLA-DR as an entry receptor for HPAI clade 2.3.4.4b H5 influenza.

## Breadth of HLA-DR specificity across H5 phylogeny

To determine the evolutionary distribution of MHC-II binding among H5 influenza viruses, we examined representative hemagglutinins spanning major clades of the H5 phylogeny, including historical clades and multiple contemporary clade 2.3.4.4b viruses (Fig. 3a). We stained neuraminidase treated, pooled human PBMCs with H5 variant tetramers and did not detect HLA-DR-dependent binding in historical H5 viruses representing clades 1, 2.1.3.2, 2.3.2.1, and 2.3.4 lineages (Fig. 3b). In contrast, all clade 2.3.4.4b viruses tested exhibited substantial HLA-DR-dependent binding, despite their diverse avian and mammalian hosts and geographic origins. Furthermore, we detected significant HLA-DR-dependent binding of the HA of A/Red Knot/Delaware Bay/2020 (H5N3), a highly diverged lineage of low pathogenic avian influenza viruses endemic to North America.

**Figure 3.**
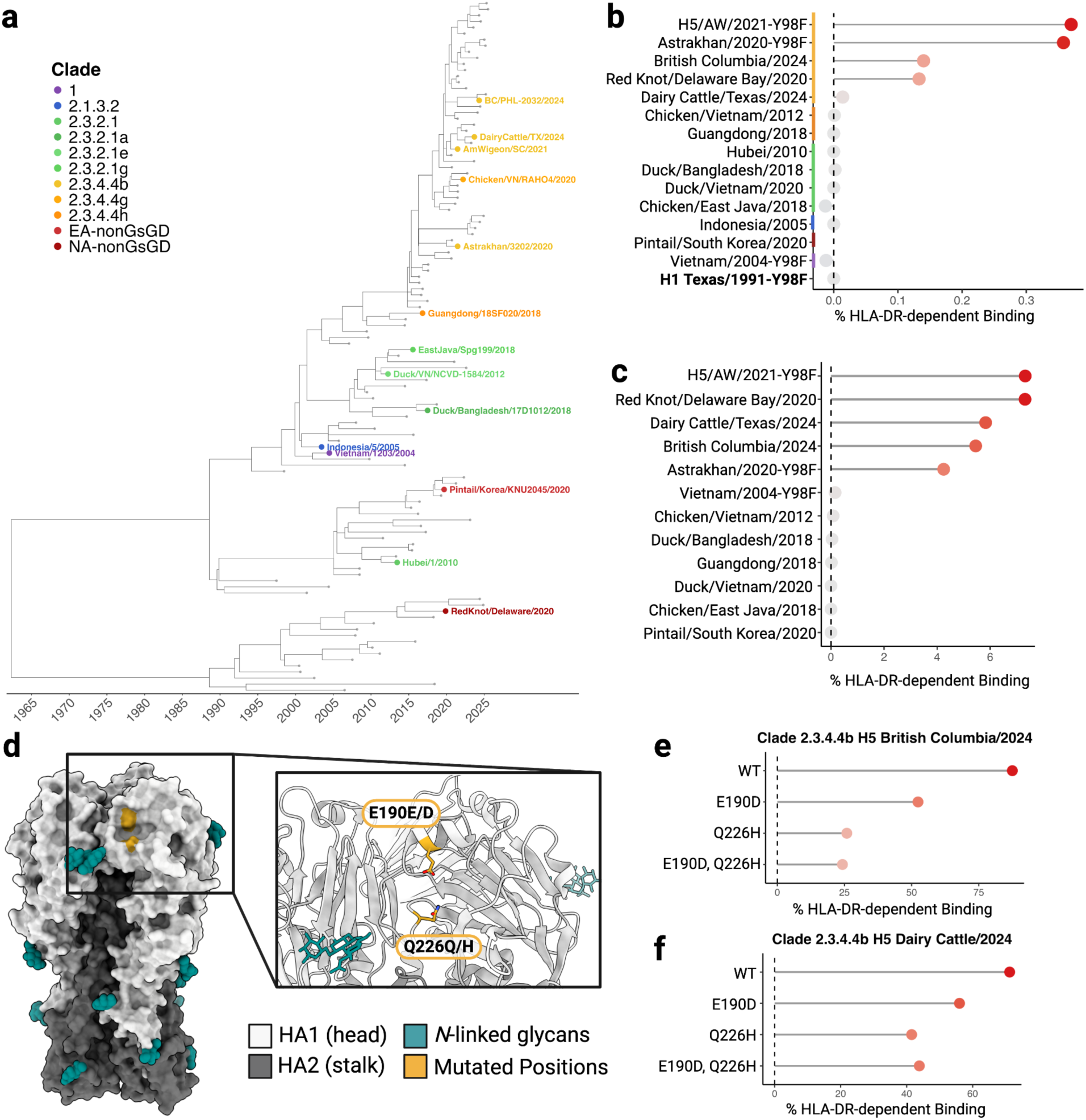
Breadth of MHC-II binding and effect of mammalian acquired mutations. **a,** Maximum-likelihood hemagglutinin phylogeny of H5Nx HA sequences (n=114 shown) with time-scaled axis. Focal strains (n=14, colored by clade) are highlighted against background sequences (grey). **b,** Lollipop graph of the HLA-DR-dependent binding of representatives from various H5 HA clades in pooled human PBMCs with colored bar to indicate H5 clade as indicated in **a**. **c,** Lollipop graph of the HLA-DR-dependent binding of representatives from various H5 HA clades in HEK293T cells transiently transfected with HLA-DR. **d,** Surface representation of prototypic H5 strain A/Jiangsu/NJ210/2023 (PDB 9MQ2^26^) with mammalian adaptation mutations highlighted in yellow. The inset shows the region of interest in ribbon representation with polymorphic residues shown as yellow sticks. **e,** Lollipop graph of the HLA-DR-dependent binding of wildtype clade 2.3.4.4b H5/British Columbia/2024 strain and mutants. Location of mutations in H3 numbering. **f,** Lollipop graph of the HLA-DR-dependent binding of wildtype clade 2.3.4.4b H5/Dairy Cattle/Texas/2024 strain and mutants.

To validate these findings, we stained HEK293T cells transfected with HLA-DR with the H5 variant tetramers. The staining data were consistent with our findings in primary cells, with HLA-DR-dependent binding enriched in clade 2.3.4.4b viruses, with the highest levels observed for American Wigeon/2021, whereas historical H5 viruses had near-background levels of binding (Fig. 3c). Together, these data indicate that HLA-DR binding is not a widespread feature of H5 hemagglutinins but rather, a new development in contemporary clade 2.3.4.4b viruses.

## Mammalian adaptation HA mutations reduce HLA-DR binding

Certain HA gene mutations have been well-characterized for facilitating the transition in influenza viruses from avian hosts toward humans and other mammalian hosts^8^. Specifically, E190D and Q226L (H3 numbering system) have been shown to alter HA sialic acid binding from the *α*-2,3 preference seen in many avian influenzas to an *α*-2,6 binding preference facilitating mammalian infection^9,10^. Notably, the E190D and Q226H polymorphisms were observed in a recent human H5 influenza infection case in British Columbia (Fig. 3d) with a frequency of detection in clinical samples at 28% and 35%, respectively, where an H5 clade 2.3.4.4b strain underwent prolonged replication in a patient, raising concern for viral adaptation for human specific receptors^11,12^. Interestingly, the effect of these mutations is strain-dependent across clade 2.3.4.4b H5 strains. Studies characterizing their effect on a British Columbia isolate confirm these mutations do not confer *α*-2,6 SA binding^11^ but rather reduce *α*-2,3 SA binding and viral fitness^13,14^. Similar mutations in dairy cattle isolates indicate they confer *α*-2,6 SA binding^15^. Given the reduced SA-dependent binding by the British Columbia viruses with these mutations, we hypothesized that selection for these HA sequence variants in a human host might be due to enhanced HA binding to HLA-DR and a corresponding increase in viral fitness. However, in neuraminidase-treated HEK293T cells transfected with HLA-DR, binding of H5/British Columbia/2021 HA variants containing E190D and Q226H substitutions was decreased compared to the wildtype HA (Fig. 3e). The same reduction in E190D and Q226H mutant HA tetramer binding to cells was also observed with a second clade 2.3.4.4b HA, H5/DairyCattle/2024 (Fig. 3f). These findings indicate that the selection for E190D and Q226H in this human infection was not due to a viral fitness increase resulting from increased HLA-DR binding.

## Binding of H5 clade 2.3.4.4b HA to primary human cells varies according to an individual’s HLA-DRB genotype

The gene encoding the human HLA-DR beta chain is highly polymorphic^16–18^ and the distribution of alleles across human population groups varies greatly^19^. To investigate whether HLA-DR encoded by certain alleles bound clade 2.3.4.4b H5 HAs more than others, we analyzed H5/AW/2021-binding B cell frequencies using primary splenic B cells from a collection of samples from deceased human organ donors for whom HLA locus genotypes were available (Fig 4a.). We observed substantial inter-donor differences in H5/AW/2021-Y98F tetramer-positive cell frequencies, ranging from 0.45% to 52.8% (Fig. 4b & 4c), while donors with the same DRB1 genotype showed some concordance in binding frequency, suggesting that DRB1 allelic variation may contribute to this inter-individual heterogeneity. However, H5/AW/2021-Y98F binding was also strongly associated with MHC-II surface abundance (Spearman’s ρ = 0.87; odds ratio = 3.04 per s.d. increase in MHC-II mean fluorescence intensity; P = 8.1 × 10^−24^; Fig. 4d). Thus, variation in MHC-II expression is likely to contribute substantially to differences in H5 binding. Because MHC-II expression is itself allele dependent^20^, allele-associated differences in binding may reflect both variation in HLA-DR receptor abundance and expression-independent differences in H5 HA binding due to HLA-DR beta chain protein sequence changes. We used ridge regression to estimate the total association of individual DRB1 alleles with H5 binding, deliberately omitting MHC-II expression from the model (Fig. 4e). DRB1*04:07, DRB1*03:01 and DRB1*04:04 showed the strongest positive associations with binding, whereas DRB1*07:04, DRB1*09:01, and DRB1*07:01 were associated with lower binding. These estimates capture the combined effects of allele-dependent expression, sequence, and other allele-specific properties on H5 binding. These findings indicate that both the HLA-DR allelic expression, protein sequence variation, and potentially other DRB1 allele-associated factors may impact the binding of clade 2.3.4.4b H5/AW/2021-Y98F HA to primary cells.

**Figure 4.**
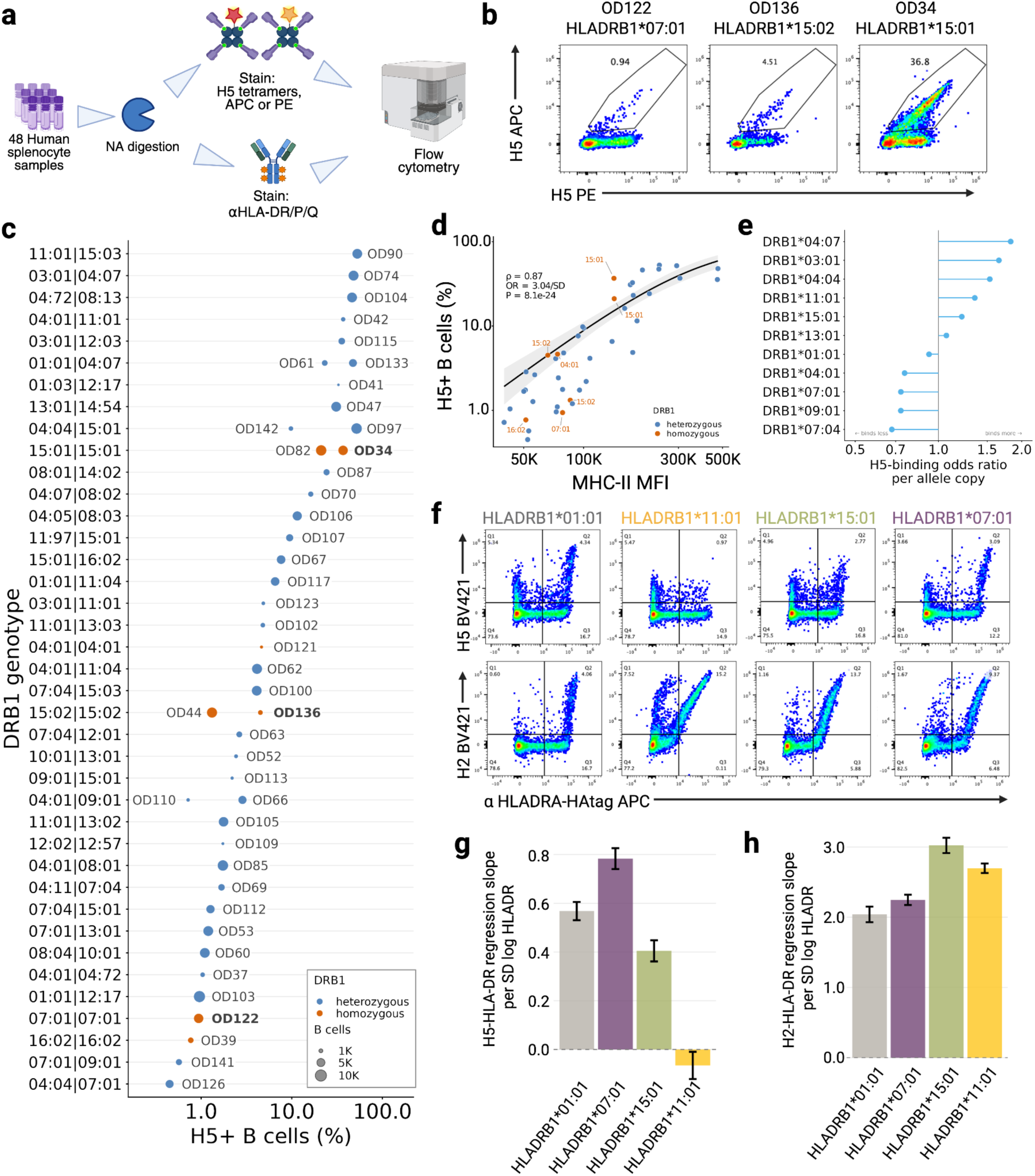
Human HLA-DR Allele specific affinity of clade 2.3.4.4b. **a**, Experimental design: splenocytes from each donor were split and stained in parallel with the H5aw-Y98F tetramers or a pan-HLA-class-II antibody (total HLA-DR/DQ/DP surface MFI); DRB1 was genotyped at two-field resolution. n = 44 biologically independent donors. **b**, Representative flow cytometry (H5 versus HLA-DR) for donors OD136, OD122 and OD34 (shown bold in c). Pre-gated on Live B cells, the gate shows H5aw-Y98F positive cells. **c**, H5⁺ fraction of CD19⁺ B cells for each donor, grouped by DRB1 genotype. Each point is one donor; point size is proportional to the number of CD19⁺ B cells analysed; color denotes zygosity (orange, homozygous; blue, heterozygous). **d**, H5⁺ fraction versus HLA-class-II MFI. Line, beta-binomial generalized linear mixed model (glmmTMB) fit of H5⁺/H5⁻ CD19⁺ counts on standardized log₁₀(MFI); shaded band, 95% confidence interval. Odds ratio = 3.04 (95% CI 2.45–3.77) per s.d. of log-expression; two-sided Wald P = 8.2 × 10⁻²⁴; Spearman ρ = 0.87. Points coloured by zygosity as in c; homozygous donors labelled with their allele. **e,** Total association of each DRB1 allele with H5 binding. All two-field alleles carried on ≥ 3 chromosomes (3–8 chromosomes each) were fitted jointly in a ridge-penalized logistic model of H5-positivity; points are the odds ratio per allele copy (log scale), referenced to 1 (vertical line). Expression is deliberately omitted, so values are total associations rather than expression-adjusted effects. **f,** Representative FC plots of live HEK293T cells stained with H5/AW/2021-Y98F tetramer (top row) or H2/Japan/1957-Y98F tetramer (bottom row) with tetramer signal plotted against HLADRA-HAtag signal colored by HLA-DRB1 allele. **g,** Bar plot of the HLA-DR dose-response slopes by allele for H5/AW/2021-Y98F. Bars represent estimated regression slopes per SD log HLA-DR expression from generalized linear mixed models of tetramer positivity; error bars show 95% confidence intervals. **h,** Bar plot of the HLA-DR dose-response slopes by allele for H2/Japan/1957-Y98F. Bars represent estimated regression slopes (per SD log HLA-DR expression) from generalized linear mixed models of tetramer positivity; error bars show 95% confidence intervals.

To further assess whether H5- and H2-derived HA tetramers showed differential binding among HLA-DR alleles, we used HEK-293T cells transiently transfected with HLA-DRA*01:01 (encoding the alpha subunit of the HLA-DR heterodimer) and one of four HLA-DRB1 alleles selected from the primary-cell analysis. We pre-treated the cells with neuraminidase to digest surface sialic acid, either blocked cells with HLA-DR antibody or left them unblocked, then stained the cells with H5/AW/2021-Y98F or H2/Japan/1957-Y98F tetramers and assayed binding by flow cytometry (Fig. 4f). We used generalized linear mixed models to test whether HLA-DR allele identity modulates the relationship between cell-surface HLA-DR expression and tetramer positivity in HEK293T transfectants. We analyzed binding for four HLA-DR alleles tested for two HA tetramers in two technical replicates for each, gated into HLA-DR+/− and tetramer+/− quadrants (Fig. 4f). For H5aw-Y98F tetramers, HLA-DR expression was a significant predictor of tetramer positivity, but the strength of this relationship differed dramatically by allele (*χ*² = [from LRT], p < 0.0001). HLA-DRB1*07:01 demonstrated the steepest expression-dependent binding response curve (slope = 0.783 per SD log HLA-DR; 95% CI: 0.740–0.826), followed by DRB1*01:01 (slope = 0.568; 95% CI: 0.531–0.605). HLA-DRB1*15:01 showed reduced correlation with expression (slope = 0.405; 95% CI: 0.361–0.448), while HLA-DRB1*11:01 exhibited minimal H5 tetramer binding signal despite HLA-DR expression comparable to the other allelic variants (slope = −0.066; 95% CI: −0.122 to −0.010) (Fig. 4g). All pairwise allele-by-slope comparisons were highly significant (Tukey-adjusted p < 0.0001 for all six comparisons). In contrast, H2/Japan/1957-Y98F tetramer binding showed a fundamentally different allele-response pattern. All four alleles demonstrated steep HLA-DR expression-dependent binding response relationships, with slopes ranging from 2.04 to 3.02 per SD log HLA-DR — roughly 3 to 50-fold steeper than those observed for H5/AW/2021-Y98F (Fig. 4h). Moreover, the allele ranking was completely reversed: HLA-DRB1*15:01 showed the steepest slope (3.02; 95% CI: 2.91–3.13), followed by HLA-DRB1*11:01 (2.70; 95% CI: 2.63–2.76), HLA-DRB1*07:01 (2.25; 95% CI: 2.17–2.32), and HLA-DRB1*01:01 (2.04; 95% CI: 1.93–2.15). All pairwise slope differences remained statistically significant after Tukey correction (p range: 0.0117 to <0.0001). Notably, HLA-DRB1*11:01, which was entirely non-responsive to HLA-DR expression for H5/AW/2021-Y98F tetramers, became one of the most H2-responsive alleles, suggesting that this allele’s interaction with HLA-DR is highly HA-variant-dependent. Together these findings suggest that a combination of allele-specific expression patterns as well as differences in binding affinity for allele-encoded HLA-DR proteins can contribute to binding by clade 2.3.4.4b H5 and H2 HAs.

## Cross-reactive human monoclonals target HLA-DR binding epitopes

HA-specific B cells often show substantial cross-reactivity to antigenically distinct influenza strains, including viruses to which the host has never been exposed^21–24^. Consistent with this, individuals naïve to clade 2.3.4.4b H5 can possess serum antibodies capable of binding clade 2.3.4.4b H5 HA^25,26^. Studies have shown that residues implicated in HLA-DR interactions lie adjacent to the canonical sialic acid receptor binding site pocket in the H2 HA^5^, suggesting that antibodies targeting this region to neutralize sialic acid-dependent entry might also interfere with HLA-DR-mediated entry. We therefore hypothesized that pre-existing cross-reactive anti-HA antibodies that bind clade 2.3.4.4b H5 HA might include examples that specifically inhibit HLA-DR-dependent viral entry, with or without an effect on binding to sialic acid.

To test this, we isolated H5 HA-binding B cells from splenocyte, mesenteric and mediastinal lymph node suspensions of deceased organ donors (n=11) using a DNA-tagged antigen panel including seven influenza HAs, including clade 2.3.4.4b H5/AW/2021-Y98F HA (indicated in Table S1), similar to a single-cell transcriptomic experimental strategy for antigen-specific B cell analysis we have previously reported for SARS-CoV-2-specific B cells^27^ (Fig. 5a). We pre-treated B cells with neuraminidase and pre-incubated them with an anti-MHC-II antibody prior to HA tetramer staining to enable specific sorting of BCR-mediated H5 tetramer binding B cells. We expressed 18 unique H5-binding native paired monoclonal antibodies (mAbs) from the single-cell BCR sequences, with H5 specificity inferred from the DNA tag counts. H5 specificity was confirmed by mAb expression and testing of H5 antigen binding in a multiplexed Luminex assay (Fig. 5b). These mAbs showed diverse VH gene usage and somatic hypermutation patterns consistent with antigen-experienced B cells, suggesting that the H5 binders were derived from cross-reactive BCRs stimulated by prior influenza exposure, likely from H1 or H2 virus exposure or H1 vaccination. This was supported by testing the 18 H5-binding mAbs for binding to a panel of other HAs in a Luminex assay (Fig. 5b) which revealed substantial diversity of binding breadth and HA cross-reactivity. Additionally, we screened the mAbs against a panel of H5 variants spanning additional major clades across the H5 phylogeny, including historical clades and an expanded number of clade 2.3.4.4b representatives (n=14, including H5 headless) (Fig. 5c). The mAbs displayed a range of binding breadths, with some only recognizing two H5 variants and others binding all except the H5 “headless” stalk-only construct. Additionally, five mAbs demonstrated neutralizing activity in a multicycle neutralization assay using an H5/Dairy Cattle/2024 conditionally replicative reporter virus where the HA and NA of H5/Dairy Cattle/2024 are expressed in the PR8 background but the PB1 gene segment is replaced with GFP in a cells that do not express HLA-DR (Fig 5b). Of the neutralizing antibodies, two show specificity for the stalk suggesting they act through inhibition of the conformational change required for fusion while remaining three likely target the sialic acid binding domain.

**Figure 5.**
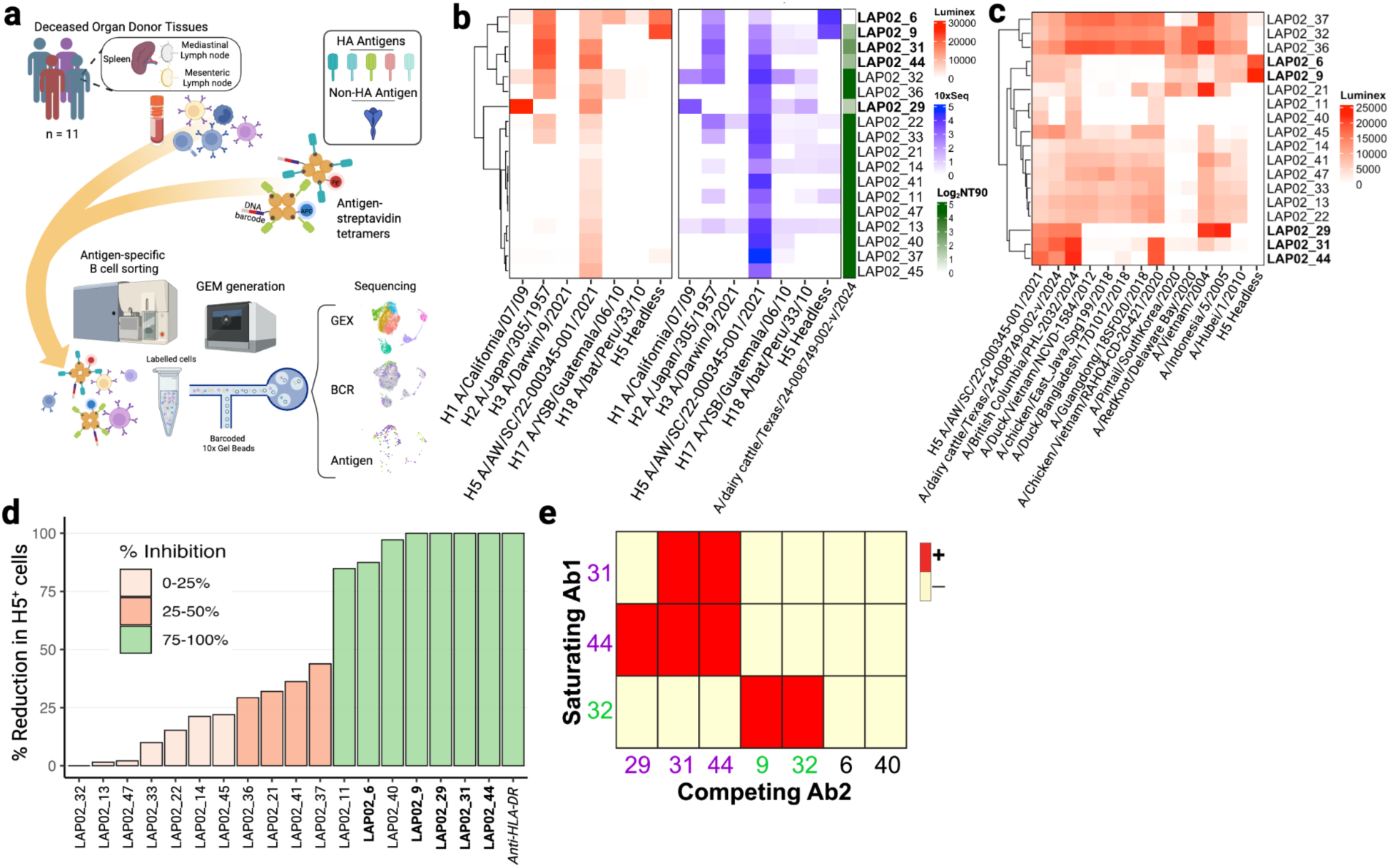
Isolation and characterization of clade 2.3.4.4b H5-specific monoclonals from naive individuals. **a,** Schematic of experimental workflow for the sorting of HA-specific B cells from deceased organ donor (n=11) single cell suspension of lymphoid tissues and recovery of transcriptomic, B cell receptor and antigen binding score via 10x Genomics 5’ single cell RNAseq. **b,** Heatmap of 10x antigen DNA counts (red) and Luminex binding scores (blue) for various HAs (n=7) and log_2_NT90 activity against pseudotyped H5/Dairy Cattle/2024 conditionally replicative reporter virus (green) for monoclonal antibodies (n=18) isolated from deceased organ donor tissues. Antibodies with neutralizing activity in bold. **c,** Heatmap of Luminex binding scores for extended H5 HA panel (n=14). **d,** Bar graph of percent reduction in H5AW tetramer positive, HLA-DR+ HEK293T cells. Control blocking antibody in italics. **e,** Shown is a hierarchically clustered heatmap for wild-type H5AW epitope binning results for three saturating mAbs (y axis) with seven competing mAbs (x axis). Binding inhibition is shown in red and normal binding is in yellow. Two epitope bins are shown by different mAb name colors

To test whether mAbs could block HA binding to human HLA-DR, we performed binding assays using clade 2.3.4.4b H5/AW/2021-Y98F HA tetramers pre-incubated with individual mAbs at saturating concentrations (30 μg/mL). HEK-293T cells transiently expressing HLA-DR were neuraminidase-treated then incubated with mAb-HA tetramer complexes in the presence of 30 µg/mL of mAb. The mAbs screened showed substantial heterogeneity in their blocking of HLA-DR-dependent binding by H5/AW/2021-Y98F HA in the neuraminidase-treated HLA-DR positive cells (Fig. 5d). Four mAbs (LAP02_29, LAP02_31, LAP02_44, and LAP02_9) achieved complete inhibition of HLA-DR-mediated H5/AW/2021-Y98F tetramer binding (100% inhibition), comparable to the positive control anti-HLA-DR monoclonal (100% inhibition). Additional mAbs demonstrated intermediate blocking: LAP02_40 (97.1% inhibition), LAP02_6 (87.4% inhibition), and LAP02_11 (84.8% inhibition). The remaining 11 mAbs showed reduced blocking capacity, ranging from 43.8% (LAP02_37) to <5% (LAP02_13, LAP02_32, and LAP02_47), suggesting that only a subset of cross reactive human antibodies to H5 HA bind epitopes at or near the HLA-DR binding site. All three neutralizing monoclonals likely targeting the SA-binding domain showed complete HLA-DR-dependent binding inhibition (Fig. 5d, bolded mAbs). Notably, two monoclonals that did not neutralize H5 for cells expressing SA, LAP02_40 and LAP02_11 also demonstrated high inhibition (97.1% and 84.8% respectively) of HLA-DR-dependent binding, suggesting that the HLA-DR binding site only partially overlaps with the SA binding domain.

To further understand the interaction between these mAbs and HA, we grouped the mAbs into epitope bins based on competitive binding measured by biolayer interferometry. Epitope binning revealed overlapping epitopes for the neutralizing antibodies, with a separate competition group and epitope bin for the non-neutralizing, HLA-DR blocking antibody, LAP02_40 (Fig. 5f). These results demonstrate that antibodies targeting the HLA-DR-binding regions of H5 HA are not rare in humans who have had routine influenza virus or vaccine exposure. Such antibodies and memory B cell clones could have the potential to provide some level of protection in the face of spillover not only through traditional SA-dependent attachment but also by disrupting association with the secondary receptor HLA-DR.

## Discussion

Clade 2.3.4.4b H5 influenza viruses have had remarkable success in infecting a wider than usual range of avian hosts, spreading globally, and recently spilling over into a wide range of mammalian species. Sustained mammal-to-mammal transmission is documented in multiple species including minks in European fur farms, South American marine mammals, and US dairy cattle. Spillover of infection into humans is clearly documented, although without infectious human-to-human transmission^8,28,29^. Our results indicate that clade 2.3.4.4b H5 influenza viruses can now be grouped with the small number of prior examples of influenza types that can use HLA-DR as a second receptor on host cells, in addition to traditional sialic acid. Contemporary clade 2.3.4.4b H5N1 viruses retain broad tissue tropism characteristic of historical H5N1 lineages^30–32^, but exhibit context-dependent disease severity differences between host species. Clade 2.3.4.4b H5N1 isolates demonstrate increased systemic dissemination and heightened pathogenic potential, particularly in cats, mice, and some other mammalian species^33–35^. The enhanced pathogenicity appears driven by factors unrelated to hemagglutinin SA receptor specificity, suggesting that contemporary clades maintain ancestral SA tropism^32^ while exhibiting differential disease manifestations in spillover species. Studies conducted in mice showed enhanced neurotropism of clade 2.3.4.4b H5 bovine isolates compared to prototypic clade 1 H5 strain, A/Vietnam/2004^36^. Whether HLA-DR binding by clade 2.3.4.4b H5 influenza viruses may play a role in host species range, cellular tropism or disease manifestations was not tested *in vivo* in this study, but we note that HLA-DR expression on immune privileged cells with ability to travel throughout the body could contribute to the trafficking of virus to the CNS as well as other tissues. Given the heterogeneity of H5-HLA-DR binding in humans, species-specific MHC-II binding could potentially be a factor contributing to variable pathogenicity and host susceptibility. Dual receptor engagement by human respiratory viruses, including the use of sialic acid in combination with cell-surface proteins, has been identified in other viral species, such as the endemic coronaviruses HCoV-OC43 and HKU1, so the belated recognition of similar biology in influenza emphasizes recurrent patterns of adaptation to the human host between viruses that infect in similar tissue environments.

We find that HLA-DR binding by H5 can contribute to cell entry by the virus, with the caveat that it is difficult to exclude a cooperative role with some residual sialic acid binding, even after thorough neuraminidase digestion. The neuraminidase used in this study has lower activity for destruction of *α*-2,6 sialic acid, compared to *α*-2,3 sialic acid, therefore, residual sialic acid in our treated cells is more likely to be the *α*-2,6 variety. It is interesting to note that a recent structure of an H5 HA (A/Texas/37/2024) showed its sialic acid binding site to be occupied by sialic acid attached to the N169 glycan of an adjacent protomer^37^. This finding raised the question of how such auto-glycan binding competes with binding to host-cell receptors. Our results suggest that HLA-DR may provide an alternative entry route in these scenarios.

An important result from the human primary cells in this analysis is that an individual’s HLA-DR genotype affects the extent of HLA-DR-dependent binding of their cells by the clade 2.3.4.4b HPAI H5 HA. Due to the correlation between allelic variants at this locus, and the expression of the HLA-DR antigen, the inter-individual H5 HA binding differences could be related to either HLA-DR expression, or protein sequence variation, or both. Our data suggest that expression differences between alleles are likely to account for most of the differences in H5 binding to primary B cells. An implication of the HLA-DR allelic differences in H5 HA binding is that different human individuals or populations, based on their HLA-DR genotypes, could have differential susceptibility to infection or disease caused by clade 2.3.4.4b H5 HPAI viruses. We do not have evidence for or against these possibilities *in vivo* in the current results, but examination of the HLA-DR genotypes in known human infection cases, and correlation with disease phenotypes and severity will be important for further evaluation of the potential role for HLA-DR in H5 infections in humans. The potential role of HLA-DR binding by clade 2.3.4.4b H5 HA in viral pathogenicity, systemic dissemination, and tissue tropism also warrants further research in animal models.

Notably, despite the low number of confirmed human cases of infection by clade 2.3.4.4b H5 influenza, a fraction of unexposed healthy individuals are nonetheless seropositive for clade 2.3.4.4b H5-binding antibodies and even neutralizing antibodies, and many individuals have memory B cells that can recognize H5 HA^26,38,39^. Studies suggest these antibodies and memory B cells are most likely derived from B cell clones stimulated by other group I influenza viruses or vaccine antigens, particularly H1 influenza viruses, or, in older individuals who were alive during the 1957-68 pandemic, H2 influenza virus^38^. Our analysis of H5-binding human mAbs was facilitated by altering our B cell sorting protocols to include blocking clade 2.3.4.4b H5 HA binding to HLA-DR to enrich for truly specific H5-binding B cells that recognize H5 HA via their BCR. We identified human mAbs that specifically block H5 binding to HLA-DR, as well as other examples that neutralize H5 infection of cells in the presence of sialic acid and HLA-DR, showing that human humoral immune responses can target these binding activities independently, but also indicating that binding of a single antibody to its epitope can interfere with both receptor binding interactions. These results are consistent with the two receptor binding sites being close to each other or overlapping in the RBS region of the HA head. Consistent with pre-immune studies in the ferret infection model^40^, the existence of these cross-reactive antibodies and memory B cells could potentially contribute to some greater protection against infection or disease caused by clade 2.3.4.4b H5 HPAI, as has been previously inferred from birth-year analyses of case mortality rates in prior H5 influenza outbreaks^41^.

Limitations of the current study include biosafety-related constraints on our ability to test actual infectious clade 2.3.4.4b H5 influenza viruses and their sequence variants, as well as the likely presence of some residual sialic acid on cells treated with neuraminidase digestion. Experiments testing binding of clade 2.3.4.4b H5 HA to different HLA-DR antigens relied on cell-surface expression of HLA-DR, either by primary cells or following transfection of HEK293T cells. Despite testing many experimental variations, as with other prior publications describing H2, or bat influenza H17 and H18 HA interactions with HLA-DR^3,5,42^, we did not detect direct binding between purified soluble clade 2.3.4.4b H5 or H2 HA proteins and soluble HLA-DR antigens loaded with CLIP in either the monomeric or tetramer form using biolayer interferometry. We suspect that this reflects a low affinity of the binding interaction, requiring the greater avidity of multivalent binding at the cell surface, but we cannot exclude other potential factors such as additional binding partners needed for complex formation (for example, invariant chain or other peptides associated with HLA-DR). Our findings motivate further evaluation of the functional role of HA-HLA-DR interactions in viral infections *in vivo*, or during vaccination with H5 HAs, as well as incorporation of data related to these receptor interactions in epidemiological modeling and spillover prediction efforts for H5 influenza.

## Acknowledgements

We would like to thank the study blood donor participants, and deceased organ donors and their families, for their generosity in making the study possible. We also acknowledge Dr. Beau Kelly, Eddie Ramos, Shelby Shortridge, and Kyle Biggs, as well as the entire team at Sierra Donor Services.

## Funding

This work was supported by NIH/NIAID CEIRR contract 75N93021C00015 (S.E.H., S.D.B.), NIH 1U54CA260517 (S.D.B.), HIPC grant U19AI057266 (S.D.B.), P01 grant 5P01AI153559 (S.D.B.), an endowment to S.D.B. from the David Crown Foundation; Early Postdoc Mobility Fellowship Stipend from the Swiss (O.F.W.); National Institutes of Health NRSA T32 T32OD011121 (T.P.).

## Author Contributions

Conceptualization: S.D.B, T.P., A.M., O.W.

Methodology: T.P., A.M., O.W., J.T.O., S.H.L, R.K.A.

Investigation: T.P., A.M., O.W., J.T.O., S.A.J., J.A., X.H., R.A.H., J-Y.L., B.L., J.S., K.R.

Sample collection: C.N., E.H., T.P., A.M., O.W., J-Y.L., B.L., X.H.

Formal analysis: T.P., A.M., O.W., S.A.J.,

Visualization: T.P., A.M., O.W., S.A.J.,

Writing—original draft: T.P., S.D.B, A.M.

Writing—review and editing: all authors

Supervision: S.D.B, S.E.H.

Funding acquisition: S.D.B., S.E.H.

## Competing interests

S.D.B. has consulted for Regeneron, Sanofi, Novartis, Genentech, Pfizer, Visterra, and Otsuka on topics unrelated to the research presented here; owns stock in AbCellera Biologics; and is a scientific cofounder of Immunera, Inc.; S.E.H reports receiving consulting fees from Sanofi, Pfizer, Lumen, Novavax, and Merck.

## Methods

### Organ donors tissue samples

Spleen, whole blood, mediastinal lymph nodes, and mesenteric lymph nodes specimens from deceased organ donors were collected from May 2021 to Jan 2025. All deceased donors were clinically managed according to established organ donation protocols, which may have included administration of 5g of methylprednisolone. Following the Uniform Anatomical Gift Act (UAGA), deceased organ donors had documentation of separate authorization for donation and research, respectively. Authorization was provided either as legal next-of-kin, or first-person authorization (e.g., registration with the Department of Motor Vehicles, DMV). As such, the Research Committee of Sierra Donor Services, Sacramento, CA and Donor Alliance, Denver, CO approved the collection of specimens from deceased organ donors who had research authorization.

### Peripheral blood mononuclear cell samples

Leukocyte reduction system chambers from healthy blood donors used for initial testing of H5 HLA-mediated binding were obtained from the Stanford Blood Center with Stanford School of Medicine Institutional Review Board approval. Samples were collected with applicable institutional guidelines and regulations and were provided to investigators in a de-identified manner; all donors provided informed consent. LRS chambers were rinsed with PBS and the recovered cell suspension was diluted and subjected to Ficoll-based density-gradient centrifugation. Peripheral blood mononuclear cells were collected from the interface and cryopreserved in fetal bovine serum with 10% DMSO in 20 mln. cell aliquots for subsequent experiments.

### Cell culture and HLA-DR expression

HEK293T cells were cultured in DMEM supplemented with 10% fetal bovine serum and penicillin–streptomycin. For transfection experiments, cells were plated at 6×10⁵ cells per well in 6-well tissue culture plates in antibiotic-free media (DMEM supplemented with 10% FBS, without antibiotics). HEK293T cells were transfected using Lipofectamine 2000 (Life Technologies) according to the manufacturer’s protocol. For single MHC-IIexpression, cells were co-transfected with an alpha chain, either HLA-DMA1-HAtag or HLA-DRA*01:01:01-HAtag and a beta chain either HLA-DMB1-V5tag or allele-specific HLA-DRB1-V5tag constructs (HLA-DRB1*01:01, HLA-DRB1*11:01, HLA-DRB1*15:01, or HLA-DRB1*07:01), with 14 µg total DNA per well in the 6-well format. Cells were incubated with transfection complexes for 4–6 hours, after which media was replaced with complete media. For neuraminidase-treated samples, media was supplemented with neuraminidase (Clostridium perfringens, Thermo Fisher) at 0.1 U/mL overnight. Transiently transfected cells were then directly stained and assayed by FC or FACS sorted to enrich for live, V5tag+ HAtag+ double-positive cells before downstream experiments.

### Recombinant HA expression and purification

Recombinant HA proteins were expressed using pCMV-Sport6 vectors encoding full-length, codon-optimized HA sequences. HA transmembrane domains were replaced with the T4 fibritin trimerization domain, AviTag for site-specific biotinylation, and a hexahistidine affinity tag^43^. Mutations were introduced using the QuikChange II XL site-directed mutagenesis kit (Agilent). Recombinant headless H5 protein was generated as previously described through mutagenesis of a recombinant headless H1 HA plasmid^44,45^. HA plasmids and a plasmid encoding neuraminidase (NA) from A/Puerto Rico/8/1934 were co-transfected into 293F suspension cells (Thermo Fisher Scientific) at a density of 1 × 106 cells/mL at 1 μg/mL. Supernatants were collected four days later and clarified by centrifugation. Recombinant HAs were purified using Ni-NTA affinity chromatography (Qiagen) and buffer exchanged into DPBS using Amicon centrifugal filters (Millipore).

### HA tetramer preparation

HA antigens (Table S1), expressed in the Hensley lab, were biotinylated using the BirA biotin-protein ligase standard reaction kit (Avidity, BirA500) and dialyzed using a Slide-A-Lyzer MINI dialysis device (10K MWCO, Thermo Scientific 69570). Protein concentrations were quantified by BCA assay (Thermo Scientific A55864) and biotinylation was confirmed by western blot. Biotinylated HAs were tetramerized with fluorophore-conjugated streptavidin with or without DNA tags (BioLegend) at a 6:1 molar ratio (antigen to streptavidin) by adding one-fifth of the streptavidin volume to the antigen every 10 minutes on ice. Reactions were quenched with 6-fold molar excess of D-biotin (Avidity) and incubated for 20 minutes on ice. Tetramers were stored at 4 °C until use.

### Neuraminidase treatment

Neuraminidase (Clostridium perfringens, Thermo Fisher) was used to deplete α-2,3-linked sialic acids and to a lesser extent α-2,3-linked sialic acids. Cells or isolated lymphocytes were resuspended in neuraminidase (0.1–1.5 U/mL) in PBS or complete media for 30 minutes at 37 °C. Treated cells were washed twice with phosphate-buffered saline (PBS) and filtered before proceeding to staining or downstream assays. Sialic acid depletion was quantified by staining with biotinylated MAL-II (Vector Laboratories B-165-1, 1ug/mL) followed by fluorotagged streptavidin (Biolegend, 0.6ug/mL) and quantified in FC.

### Primary cell staining

Frozen human PBMCs or splenocytes were rapidly thawed at 37 °C before adding 10 mL of pre-warmed medium dropwise. Cells were centrifuged at 300 g for 10 minutes and resuspended in FACS buffer (1× PBS + 0.2% BSA) and counted. Where indicated, cells were treated with neuraminidase as described above. Untreated cells were incubated at 37C in PBS. Cells were washed twice with cell staining buffer (BioLegend 420201) before staining with anti-CD19 PE-Cy7 (BioLegend 302216) and live/dead dye (Fixable Viability Dye eFluor 780, Thermo Fisher Scientific, or Ghost780 Cytek SKU 13-0865-T100).All cells were stained for 30 minutes on ice. For antibody blocking experiments, the following monoclonal antibodies were added to the proceeding stain mix at the specified concentrations: anti-pan-MHC-II (BioLegend 361702), anti-HLA-DR (10–50 µg/mL; BioLegend 307602), anti-HLA-DQ (10 µg/mL, BioLegend 361502), and anti-CD74 (10 µg/mL,BioLegend 326802). Cells were washed once with FBS, resuspended in HA tetramers at 125 - 250 ug tetramer per 50uL reaction in CSB and incubated for 30 minutes on ice. Finally, cells are washed twice with CSB before analysis on a Cytek Aurora flow cytometer (Cytek Biosciences) for FC or BD FACS Aria for FACS.

### HEK293T Staininglow cytometry staining and antibody blocking

Cells were harvested by trypsinization,washed once with PBS, and treated with neuraminidase as described above. Untreated cells were incubated at 37C in PBS. Cells were washed twice with cell staining buffer (CSB, BioLegend 420201). Cells were stained with PE-conjugated anti-V5tag antibody (Thermo Scientific 12-6796-42) and APC-conjugated anti-HA11.1 tag antibody (BioLegend 901523). All cells were stained for 30 minutes on ice.For antibody blocking experiments, the following monoclonal antibodies were added to the proceeding stain mix at the specified concentrations: anti-pan-MHC-II (BioLegend 361702) or anti-HLA-DR (10–50 µg/mL; BioLegend 307602). Cells were washed once with FBS, resuspended in HA tetramers at 125 - 250 ug tetramer per 50uL reaction in CSB and incubated for 30 minutes on ice. For monoclonal blocking assays, tetramers were pre-incubated with monoclonals at 30ug/mL for 30 minutes prior to tetramer staining cells. Finally, cells are washed twice with CSB before analysis on a Cytek Aurora flow cytometer (Cytek Biosciences) for FC or BD FACS Aria for FACS.

### HLA-DRB Allele screen in deceased organ donors

Frozen human splenocytes from 48 organ donors were rapidly thawed at 37 °C before adding 10 mL of pre-warmed medium dropwise. Cells were centrifuged at 300 g for 10 minutes and resuspended in PBS with neuraminidase and treated as described above. After treatment cells were resuspended in CSB (BioLegend) and stained with anti-CD19 BV421, anti-CD27 PE-Fire810 (all BioLegend), and split into two halves and separately stained with tetramers or anti-HLA antibody to avoid competition effects. The first half was stained with H5aw-Y98F APC- and PE-labeled tetramers for 30 minutes on ice at 125 µg tetramer per 50 µL reaction. The second half was stained with anti-pan-MHC-II antibody APC-Fire750 (BioLegend). Cells were washed twice with CSB before analysis on a Cytek Aurora flow cytometer (Cytek Biosciences)

### Whole exome sequencing and HLA genotyping

Genomic DNA was extracted from a separate aliquot of cryopreserved cells from spleen or PBMCs from 48 deceased organ donors using Qiagen Allprep kit (Qiagen). Whole exome sequencing libraries were prepared using Illumina DNA Prep with Exome 2.5 Enrichment kit (Illumina) per manufacturer’s instructions and sequenced on NovaSeq 6000 sequencing platform (Illumina). HLA genotyping was performed using HISAT-genotype software (^46^

### GFP Reporter virus particle viral entry assays

HEK293T cells co-transfected with HLA-DRA-HAtag and HLA-DRB1-V5tag were treated with or without neuraminidase (1 U/mL, 30 minutes at 37 °C) and stained for surface markers (anti-HA11.1 tag APC, anti-V5 tag PE, ghost780) for 30 minutes on ice. FACS sorting was performed to enrich for live, V5tag+ HAtag+ transfected cells. Sorted cells were plated at 1×10⁴ cells per well in 96-well plates in complete media and inoculated with retroviral pseudotypes (RVPs) bearing influenza HA from three H5N1 strains: A/American Wigeon/South Carolina/22-000345-001/2021 (Integral Molecular, RVP-1214G), A/dairy_cattle/Texas/24-008749-003-original/2024 (Integral Molecular, RVP-1217G), or A/Vietnam/1203/2004 (clade 1 control) with GFP reporter at 2.23×10⁵ transduction units (TU) per well. For HLA-DR blocking conditions, sorted cells were pre-incubated with anti-HLA-DR (BioLegend Cat #307602, 50 µg/mL) for 30 minutes at 37 °C before RVP addition. On day 2 post-infection, cells were harvested, stained for anti-HLA-DR PE-Dazzle (BioLegend 307654), HA tag (APC), V5 tag (PE), and ghost780, and analyzed by flow cytometry. GFP+ infection rates were quantified as a percentage of live, V5+ HAtag+ transfected cells.

### Antigen-specific B cell isolation and sorting

Frozen single-cell suspensions of mesenteric or mediastinal lymph nodes and/or spleen from organ donors were thawed at 37 °C and 15 mL of pre-warmed thawing medium (RPMI + 10% FBS) was added dropwise. Cells were then centrifuged at 300 g for 10 minutes, and washed once with FACS buffer (1× PBS + 0.2% FBS). B cells were isolated using the EasySep™ Human Pan-B Cell Enrichment Kit (Stemcell, cat #19554). Samples were then washed and resuspended in 500mM Neuraminidase at 4 million cells/ml and incubated at 37 °C for 30 minutes. After washing, cells were filtered and incubated with FACS buffer, containing MHC-II blocking antibody (Biolegend) for 30 minutes on ice. Cells were then stained with antigen tetramers in Brilliant Stain Buffer (BD Horizon) with 5pM D-Biotin (Thermo Fisher Scientific, #B20656) for 1 hour on ice. Cells were then stained with 50 µL of an B cell surface marker antibody cocktail containing anti-CD19 PE-Cy7, anti-CD27 BV605, anti-CD38 AF700, anti-IgM PerCP-Cy5.5, anti-IgD PE-Dazzle594, anti-IgG FITC, (all BioLegend) in Brilliant Stain Buffer (BD Horizon) with 5pM D-Biotin (Thermo Fisher Scientific, #B20656), and Fixable Viability Dye eFluor 780 (Thermo Fisher Scientific) for 30 minutes at 4°C. Cells were washed with FACS buffer. Before sorting, samples from each tissue were pooled and resuspended in 1× PBS + 0.04% BSA at 10 million cells/mL. Three populations were FACS sorted separately into PBS + 0.04% BSA on a BD FACSAria Fusion (four-laser configuration): (1) HA-tetramer+ B cells (PE+ APC+, representing antigen-specific populations), and (2) total B cells as a negative control.

### 10x Genomics single-cell sequencing and library preparation

Sorted B cells were processed using Chromium GEM-X Single Cell 5′ v2 with Feature Barcoding technology (PN-1000263) following 10x user guide CG000330 Rev A, Rev A (March 07, 2024). The three sorted populations were loaded separately for GEM generation. The number of cDNA amplification cycles was determined by cell recovery for each population. VDJ, gene expression (GEX), and feature barcode (FB) libraries were pooled at a 1:4:1 ratio and sequenced on a NovaSeq X Plus (25B, PE150) at 150 pM final loading concentration with 1% PhiX (Novogene). Sequencing depth followed 10x user guide recommendations.

### Analysis of single-cell data

Raw sequencing data were demultiplexed and aligned to the human reference genome (GRCh38) using Cell Ranger (10x Genomics, v8.0.1). Filtered feature-barcode matrices were analyzed in R using Seurat [or Scanpy]. Quality control filtering excluded cells with <500 or >5,000 genes detected, >20% mitochondrial reads, or doublet signatures (identified via DoubletFinder or similar). VDJ libraries were processed using MiXCR v4.7.0 using the preset “10x-sc-xcr-vdj-gemx-v3”. BCR sequences were extracted and paired to their corresponding GEX profiles via barcode matching. Heavy and light chain pairs were assembled and annotated for VDJ segment usage, somatic hypermutation (SHM) rates, and isotype. Feature barcode and hashtag barcode libraries enabled assignment of antigen specificity and tissue origin to individual B cells. Antigen specificity was determined by which HA variants showed signal above background. Single B cells were selected for monoclonal antibody production if they met the following criteria: (1) live, singlet, CD19+ CD20+, (2) paired heavy and light chain VDJ sequences with productive rearrangements, (3) tetramer+ or memory marker expression, and (4) ≥1,000 transcripts detected.

### Monoclonal antibody production and validation

Codon-optimized variable regions were cloned into IgG1 heavy chain and kappa or lambda light chain plasmids (Twist Biosciences). Paired heavy and light chain plasmids were transfected at a 1:1 ratio into Expi293F suspension cells (Gibco) at a density of 3×10^6^ cells/mL using PEI MAX (Kyfora Bio) in Opti-MEM (Gibco). Four days-post transfection, cell culture supernatants were clarified by centrifugation, then purified with the KingFisher Apex system (Thermo Scientific) using Pierce High Capacity Protein A MagBeads (Thermo Scientific) according to the manufacturer’s instructions. Antibodies were eluted with IgG elution buffer (Thermo Fisher Scientific) and neutralized with 1.0 M Tris, pH 8.8, then concentrated and buffer exchanged into DPBS (Corning) using 30 kDa Amicon centrifugal filters (Millipore).

### Luminex multiplex antigen-binding assay

Full-length or headless HA proteins were coupled to MagPlex beads (Luminex) at a ratio of 0.1 nmol antigen per 10^6^ beads. For each antigen tested, 2,500 beads were added to each well of a 96-well black, clear-bottom plate. Blank beads, which were not coupled to any antigen, were included in each well as a negative control. Monoclonal antibodies were diluted to 10 μg/ml in PBS-TBN (1X PBS, 0.1% bovine serum albumin, 0.02% tween 20, and 0.05% sodium azide), then added to beads and incubated with 600 rpm shaking for one hour at room temperature. Beads were washed twice with PBS-TBN, then antigen-specific antibodies were detected with mouse anti-human IgG-PE at 2 μg/mL (SouthernBiotech) in PBS-TBN. After a 30-minute incubation, beads were washed twice with PBS-TBN, then read on the xMAP INTELLIFLEX system (Luminex). Mean fluorescence intensities (MFI) were background-corrected by subtracting the respective blank bead MFI.

### Viral neutralization assays

Recombinant influenza virus expressing the HA, mutated to contain a monobasic cleavage site, and NA of A/DairyCattle/Texas/24-008749-002-v/2024 was generated in a conditionally replicative system in which the PB1 gene is replaced by GFP^47^ and other internal gene segments are of A/Puerto Rico/8/1934, preventing replication on cells that do not express PB1. Briefly, the virus was rescued by transfecting pHW2000 plasmids along with pHH-PB1flank-eGFP and pHAGE2-EF1aInt-TMPRSS2-IRES-mCherry into a co-culture of 293T-CMV-PB1 and MDCK-SIAT1-PB1-TMPRSS2 cells using Lipofectamine 2000 (Invitrogen) in Opti-MEM (Gibco). One day post-transfection, cells were washed with DPBS (Corning) before adding neutralization assay media (NAM) comprised of Medium-199 (Gibco) with 0.01% FBS (Sigma), 0.3% bovine serum albumin (BSA; Sigma), 100 U/mL penicillin (Gibco), 100 µg/mL streptomycin (Gibco), 100 µg/mL calcium chloride (Sigma), and 25 mM HEPES (Corning). Two days later, transfection supernatants were clarified by centrifugation and expanded on MDCK-SIAT1-PB1-TMPRSS2 cells in NAM. A modified 50% tissue culture infectious dose (TCID50) assay using GFP as a readout for infection was used to determine the infectious titer of the expanded virus, using the Reed and Muench method.

Two-fold serial dilutions of monoclonal antibodies, starting at 10 μg/ml, were prepared in NAM and mixed with an equal volume of virus, diluted to yield 200 TCID50/well. Wells containing virus but no antibody and neither virus nor antibody were included on each plate as controls. After one hour, 2.5×10^4^ MDCK–SIAT1–CMV–PB1–TMPRSS2 cells were added to each well, then plates were incubated at 37°C for 42 hours to allow for viral replication. Cells were fixed with 4% paraformaldehyde (Electron Microscopy Sciences) and the mean fluorescence intensity (MFI) of GFP was determined with an EnVision microplate reader, using a 485 nm excitation wavelength and a 530 nm emission wavelength. The 90% neutralizing titers (NT90) were determined as the lowest concentration of antibody that yielded less than 10% of the average MFI of virus only wells after background correction. Antibodies that failed to achieve this were assigned an NT90 of 20 μg/ml.

### BLI epitope binning

Biotinylated purified A/American Wigeon/South Carolina/22-000345-001/2021(H5N1) recombinant HA was loaded at 50 nM onto Octet streptavidin biosensors (Sartorius, Cat#18-5020) on a Sartorius Octet RED96 biolayer interferometer. Purified IgG1s (50 nM) were first screened for binding strength and three mAbs that bound best were picked as saturating Ab1s. For epitope binning, 50 nM saturating IgG1 mAbs were bound to biosensors for 12 minutes before 25 nM competing Ab2 IgG1 mAbs were bound for 10 minutes. Fractional signals were calculated relative to the maximal possible signal obtained if buffer, not saturating Ab1, was presented before Ab2, and these values were used to determine if inhibited or normal binding was present.

### Statistical analysis

P-values were determined using two-sided non-parametric methods. Paired comparisons (e.g., untreated vs. neuraminidase-treated within donors) were analyzed using the Wilcoxon signed-rank test. Unpaired multi-group comparisons were analyzed using Kruskal-Wallis test with post-hoc Dunn correction for multiple comparisons. Correlations between HLA-DR expression and tetramer binding were assessed using Spearman’s rank correlation. For RVP entry assays, statistical analysis of % GFP+ cells used paired t-tests with Bonferroni correction when applicable. P-values <0.05 were considered statistically significant. All analyses were performed in R (v4.3.1). For the experiment using samples from multiple organ donors (Fig. 4e), carrying various HLA-DR1 alleles, we utilized ridge-penalised logistic regression to rank alleles jointly while accounting for their sparsity and for linkage disequilibrium among HLA alleles. We entered all two-field DRB1 allele dosages carried on ≥ 3 chromosomes into a single ridge-penalised logistic regression (‘glmnet’, ‘alpha = 0’, two-column binomial response), with the penalty strength λ chosen by 10-fold cross-validation (‘lambda.min’); coefficients were exponentiated to odds ratios per allele copy and displayed as a lollipop against a reference of OR = 1. Expression was deliberately excluded from this model. Fig. 4G and H, Live, singlet cells were classified into four quadrants (DR+/−, Tet+/−) via FlowJo, and each quadrant was exported separately as per-event CSV files. We fit generalized linear mixed models (GLMM, binomial family) in R via lme4::glmer() with the formula: tet_binary ∼ log_DR_z * allele + (1 | sample_id)where log_DR_z is log-scaled HLA-DR expression centered to mean 0, SD 1 to test whether HLA-DR allele modifies the relationship between log-scaled cell-surface HLA-DR expression and tetramer positivity, including fixed effects for log(HLA-DR) × allele × tetramer interaction and a random intercept for sample_id. Model convergence was verified via allFit() across six optimizers. Allele-specific slopes were estimated via emmeans::emtrends() with Tukey-corrected pairwise comparisons.

## Data availability

Antibody sequence data will be deposited in an NIH public repository upon publication. Recombinant monoclonal antibodies from this study are available from the corresponding author upon request. Raw flow cytometry data (.fcs files) and pseudoviral stocks are available from the corresponding author. Plasmid constructs encoding HLA-DR alleles and HAs are available from the corresponding author.

## Supplemental Material

**Supplemental Figure 1.**
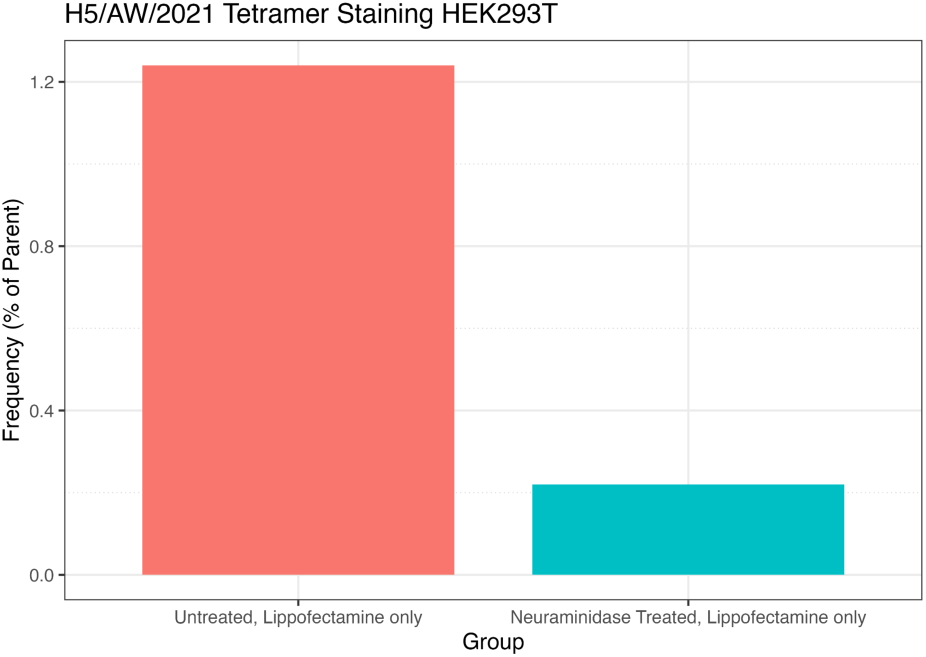
Neuraminidase treatment reduces H5/AW/2021 tetramer positive cell frequency. Bar graph of the frequency of live, tetramer positive HEK293T cells either neuraminidase treated or untreated and stained with H5/AW/2021-Y98F HA tetramer.

**Table S1.**
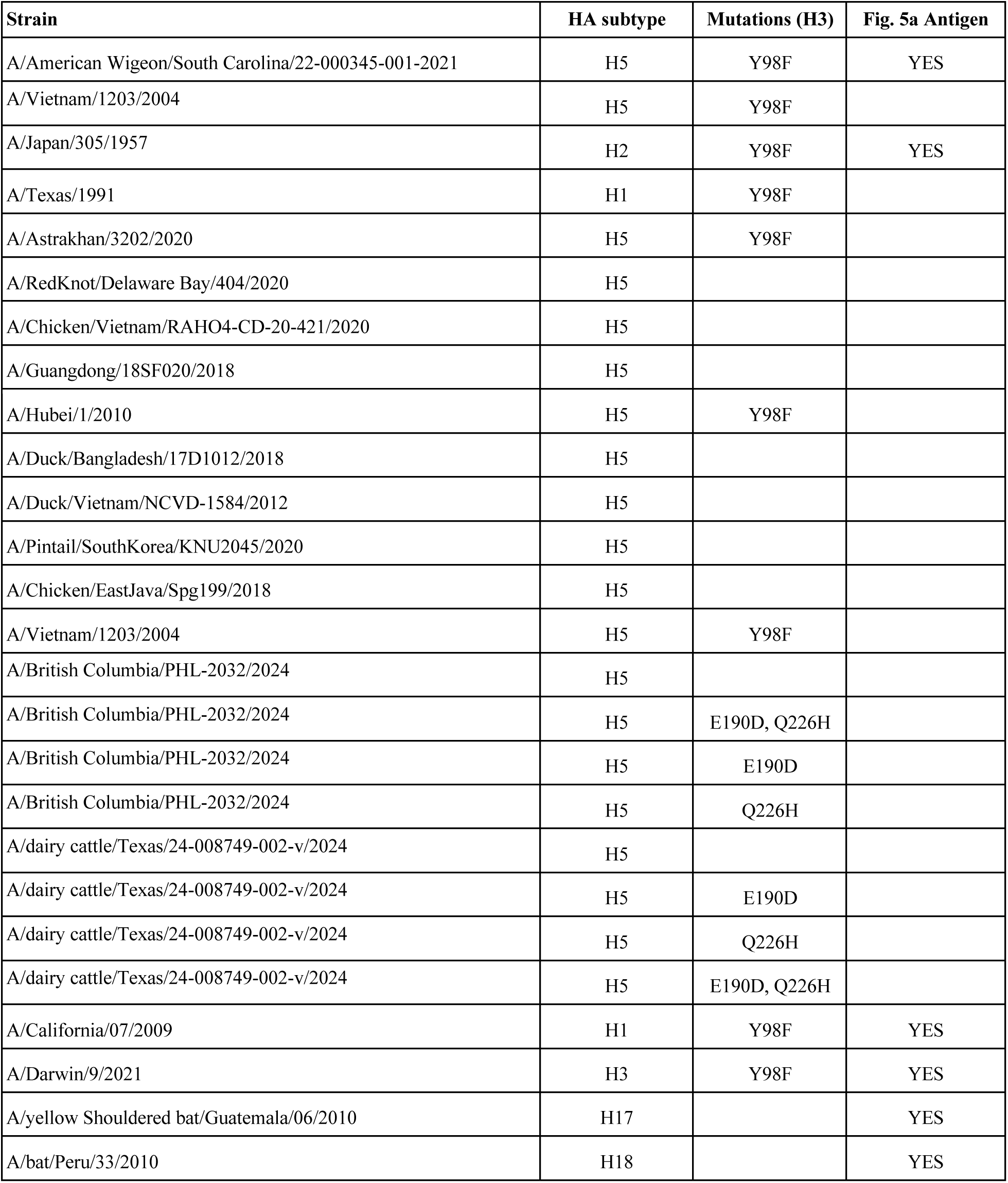
Hemagglutinin variants and metadata.

